# Evaluating methods for measuring background connectivity in slow event-related functional MRI designs

**DOI:** 10.1101/2022.12.02.518897

**Authors:** Lea E. Frank, Dagmar Zeithamova

**Author notes:** Correspondence concerning this article should be directed to Dagmar Zeithamova, 1451 Onyx St. Eugene, OR 97403.

## Abstract

Resting-state functional MRI (fMRI) is widely used for measuring functional interactions between brain regions, significantly contributing to our understanding of large-scale brain networks and brain-behavior relationships. Furthermore, idiosyncratic patterns of resting-state connections can be leveraged to identify individuals and predict individual differences in clinical symptoms, cognitive abilities, and other individual factors. Idiosyncratic connectivity patters are thought to persist across task states, suggesting task-based fMRI can be similarly leveraged for individual differences analyses. Here, we tested the degree to which functional interactions occurring in the background of a task during slow event-related fMRI parallel or differ from those captured during resting-state fMRI. We compared two approaches for removing task-evoked activity from task-based fMRI: (1) applying a low-pass filter to remove task-related frequencies in the signal, or (2) extracting residuals from a general linear model (GLM) that accounts for task-evoked responses. We found that the organization of large-scale cortical networks and individual’s idiosyncratic connectivity patterns are preserved during task-based fMRI. In contrast, individual differences in connection strength can vary more substantially between rest and task. Compared to low-pass filtering, background connectivity obtained from GLM residuals produced idiosyncratic connectivity patterns and individual differences in connection strength that more resembled rest. However, all background connectivity measures were highly similar when derived from the low-pass filtered signal or GLM residuals, indicating that both methods are suitable for measuring background connectivity. Together, our results highlight new avenues for the analysis of task-based fMRI datasets and the utility of each background connectivity method.

## 1. Introduction

Neuroimaging studies have revealed that distant brain regions can exhibit correlated neural activity, or “functional connectivity,” even in the absence of external stimulation (Clark et al., 1984; Friston, 1994; Friston et al., 1993; Shaw, 1981; Tucker et al., 1986). The most common approach to measuring functional connectivity is with resting-state functional MRI, during which spontaneous low-frequency fluctuations in brain activity are recorded in the absence of an explicit task. Resting-state functional connectivity studies have substantially contributed to our understanding of brain organization and brain-behavior relationships. A key contribution of resting-state connectivity studies has been the identification of large-scale brain networks (Power et al., 2012; D. Wang et al., 2015; Yeo et al., 2011). These functional networks align with what we know from task-based fMRI (Beckmann et al., 2005; Dosenbach et al., 2007; Fox, Corbetta, et al., 2006; Greicius et al., 2003) and are relatively stable across time and individuals (Gratton et al., 2018; Guo et al., 2012; Horien et al., 2019; Zuo et al., 2010). Although functional connectivity does not imply existence of underlying structural connections, regions that are structurally connected tend to exhibit high functional connectivity (Greicius et al., 2009; Honey et al., 2007; Passingham et al., 2002; Rykhlevskaia et al., 2008; Van Den Heuvel et al., 2009).

While network structure is relatively consistent across individuals, functional connectivity measures also contain important information about individual differences. For example, stable idiosyncratic differences exist among individuals such that a pattern of an individual’s connections may serve as their “connectivity fingerprint” (Finn et al., 2015). Furthermore, a range of studies have demonstrated the functional relevance of individual differences in connectivity, such that the strength of a specific connection or a network metric can predict individual differences in clinical (Reinen et al., 2018; Takamura & Hanakawa, 2017; Tracy & Doucet, 2015), cognitive (Finn et al., 2015; Fong et al., 2019; Rosenberg et al., 2015), or age variables (Dosenbach et al., 2010; Ferreira & Busatto, 2013; Geerligs et al., 2015; Wang et al., 2012).

While resting-state connectivity measures remain the gold standard, follow-up work has also explored the utility of connectivity measures obtained while a person performs a task. A frequently utilized functional connectivity measure in task-based designs is *background connectivity,* or coupling between regions that is observed after task-evoked activity has been controlled for (Al-Aidroos et al., 2012; Fair et al., 2007; Frank, Bowman, et al., 2019; Frank, Preston, et al., 2019; Norman-Haignere et al., 2012). Such measures can provide information about how interactions between regions are altered by different cognitive demands, for example, comparing background connectivity under different task conditions (Al-Aidroos et al., 2012; Cooper & Ritchey, 2019; Norman-Haignere et al., 2012; Tambini et al., 2017). Notably, some argue that connectivity patterns found during rest persist during different task states, with task-evoked activity comprising a modest proportion of overall connectivity structure (Cole et al., 2014; Gratton et al., 2018; Kraus et al., 2021). As Fair et al. (2007) suggested, one may not need to collect a dedicated rest scan; it may be possible to extract resting-state-like connectivity profiles from task-based fMRI, to identify network structure and measure individual differences in connectivity. With the growth of publicly available MRI data, this means researchers can get further use out of task-based fMRI datasets.

A key challenge with measuring functional connectivity in task-based designs is the potential confounding role of task-related activity. For example, two regions that are otherwise minimally functionally connected may both show task-evoked activity, such as both increasing activation in response to stimulus onset. Correlating raw activation timecourses would generate conflated connectivity estimates that do not reflect the true nature of their functional relatedness. Yet, if brain activity during a task is a roughly linear combination of spontaneous and task-evoked activations—as has been suggested (Fox, Snyder, et al., 2006; Fox & Raichle, 2007)—it may be possible to isolate resting-state-like connectivity profiles after statistically removing task-evoked activity from the observed signal (Cole et al., 2014; Fair et al., 2007; Gratton et al., 2018).

The degree to which background connectivity may approximate resting-state connectivity is, however, not yet clear. For example, Fair and colleagues (2007) showed that background connectivity estimates obtained from block-design task-based fMRI produced average connection patterns similar to those found at rest. In contrast, larger differences compared to rest were found when extracting background connectivity from a jittered event-related fMRI design. Furthermore, how “connectivity fingerprints” or individual differences in background connectivity relate to those from resting-state could not be evaluated, as the study by Fair and colleagues measured background connectivity and resting-state connectivity in different subjects.

Another question is how the correspondence between rest and background connectivity estimates may be affected by a specific method of removing task-evoked responses. For example, Fair and colleagues (2007) modeled task-evoked responses with a general linear model, using a finite impulse response (FIR) function to maximize the fit between the BOLD signal and the model (Al-Aidroos et al., 2012; Cooper & Ritchey, 2019; Duncan et al., 2014; Norman-Haignere et al., 2012). After the task-evoked signal was modelled out, background connectivity was then computed using the residual timeseries. Another approach, suited for slow event-related designs with regularly space trials, is applying a low-pass filter to remove activity fluctuations at the task frequency and leaving only lower-frequency fluctuations reflecting background activity (Frank, Bowman, et al., 2019; Frank, Preston, et al., 2019; Tambini et al., 2017). While applying a low-pass filter can be a computationally faster alternative to FIR modeling, it is unclear whether the two methods produce similar connectivity profiles and how they each compare to rest.

Here, we tested the idea that resting-state-like connectivity may be obtained from task-based fMRI after the removal of task-evoked signal, by measuring congruency between background connectivity estimates and resting state connectivity in a slow-event related fMRI design. We focused on slow event-related fMRI as it has not been formally compared to rest, and its use may be increasing due to its benefits for trial-by-trial multivariate pattern analyses (Zeithamova et al., 2017). Moreover, a slow-event related design allows for both methods of removing task-evoked activation: FIR modeling (Al-Aidroos et al., 2012; Fair et al., 2007; Norman-Haignere et al., 2012) and low-pass filtering (Frank, Bowman, et al., 2019; Frank, Preston, et al., 2019; Tambini et al., 2017). Here, we compare connectivity patterns obtained from a rest scan with background connectivity patterns obtained from low-pass filtered task-based fMRI and FIR residuals obtained from the same task-based fMRI. In addition to testing the reproduction of large-scale brain networks, we utilized subject-specific ROI-to-ROI connectivity matrices to evaluate the stability of within-subject connectivity profiles and individual differences across rest and background connectivity methods.

## 2. Methods

### 2.1. Participants

Participants were recruited from the University of Oregon and surrounding community for a larger study that included an MRI component for a subset of participants. Only data from the scanned participants are included here and the larger study will not be discussed in the present report. A total of 62 participants were scanned, six of which were excluded: four for falling asleep during the resting-state scan, one for non-compliance with study procedures, and one for not having enough data following the scrubbing procedures described below. All analyses report the final sample of 56 participants. Participants received written informed consent and were financially compensated for their time. All experimental procedures were approved by Research Compliance Services at the University of Oregon. Participants were eligible for the MRI if they were right-handed, native English speakers, had no MRI contraindications, psychiatric or neurological illnesses, and were not taking medications known to affect brain function.

### 2.2. Procedure

#### 2.2.1 Overview

In this study, participants underwent functional MRI while completing a resting-state scan and a passive viewing task. During the resting-state scan (8 minutes), participants were instructed to keep their eyes open while a fixation cross was projected onto a screen that was viewed through a mirror. Participants then completed four runs (3.67 minutes each) of task-based fMRI that consisted of passive viewing of face stimuli shown one at a time every 12 seconds (2 s stimulus, 10 s fixation inter-trial interval).

Each run started with a 4 second fixation cross, followed by a total of 18 trials (9 unique faces, each repeated twice). Participants were instructed to not make any responses during this time. In between the second and third passive viewing run, participants completed an un-scanned category learning task where they learned to sort the faces into three families. The MRI scan was part of a larger, three-visit study where participants completed a battery of tests assessing memory and other cognitive abilities. The results from task-based fMRI analyses, the categorization task, and other measures will be reported separately and are not included in the present report. Rather, here we utilize the resting-state and task-based fMRI data to address the methodological question of obtaining resting-state-like connectivity measures from task-based designs.

#### 2.2.2 fMRI Data Acquisition

Scanning was completed on a 3T Siemens Skyra at the UO Lewis Center for Neuroimaging using a 32-channel head coil. Foam padding was used around the head to minimize motion. The scanning session consisted of a localizer SCOUT sequence, an 8-minute resting-state scan, four functional 3.67-minute runs of the passive viewing task, and two anatomical scans. Functional data were acquired using a multiband gradient-echo pulse sequence (TR, 2000 ms; TE, 25 ms; flip angle, 90°; matrix size, 104×104; 72 contiguous slices oriented 15° off the AC-PC line to reduce prefrontal signal dropout; interleaved acquisition; FOV, 208 mm; voxel size: 2.0×2.0×2.0 mm; GRAPPA factor, 2; Multi-band acceleration factor, 3). Anatomical data was collected using a standard high-resolution T1-weighted MPRAGE anatomical image (TR, 2500 ms; TE, 3.43 ms; TI, 1100 ms, flip angle, 7°; matrix size, 256×256; 176 contiguous sagittal slices; FOV, 256 mm; slice thickness, 1 mm; voxel size 1.0×1.0×1.0 mm; GRAPPA factor, 2) and a custom anatomical T2 coronal image (TR, 13520 ms; TE, 88 ms; flip angle, 150°; matrix size, 512×512; 65 contiguous slices oriented perpendicularly to the main axis of the hippocampus; interleaved acquisition; FOV, 220 mm; voxel size, 0.4×0.4×2.0 mm; GRAPPA factor, 2).

#### 2.2.3 fMRI Analysis Strategy

Here, we aimed to compare two methods for removing task-related activity, low-pass filter and FIR residuals, in a slow event-related fMRI design. Each of these methods were applied to the functional data collected during the passive viewing task to remove task-evoked activity. Background connectivity was then measured and averaged across the four runs. Functional connectivity was also calculated from the resting-state scan and used as a “gold standard” to which background connectivity was compared. An overview of the preprocessing and analysis pipeline is shown in **Figure 1** and a detailed description of the steps are outlined below.

**Figure 1.**
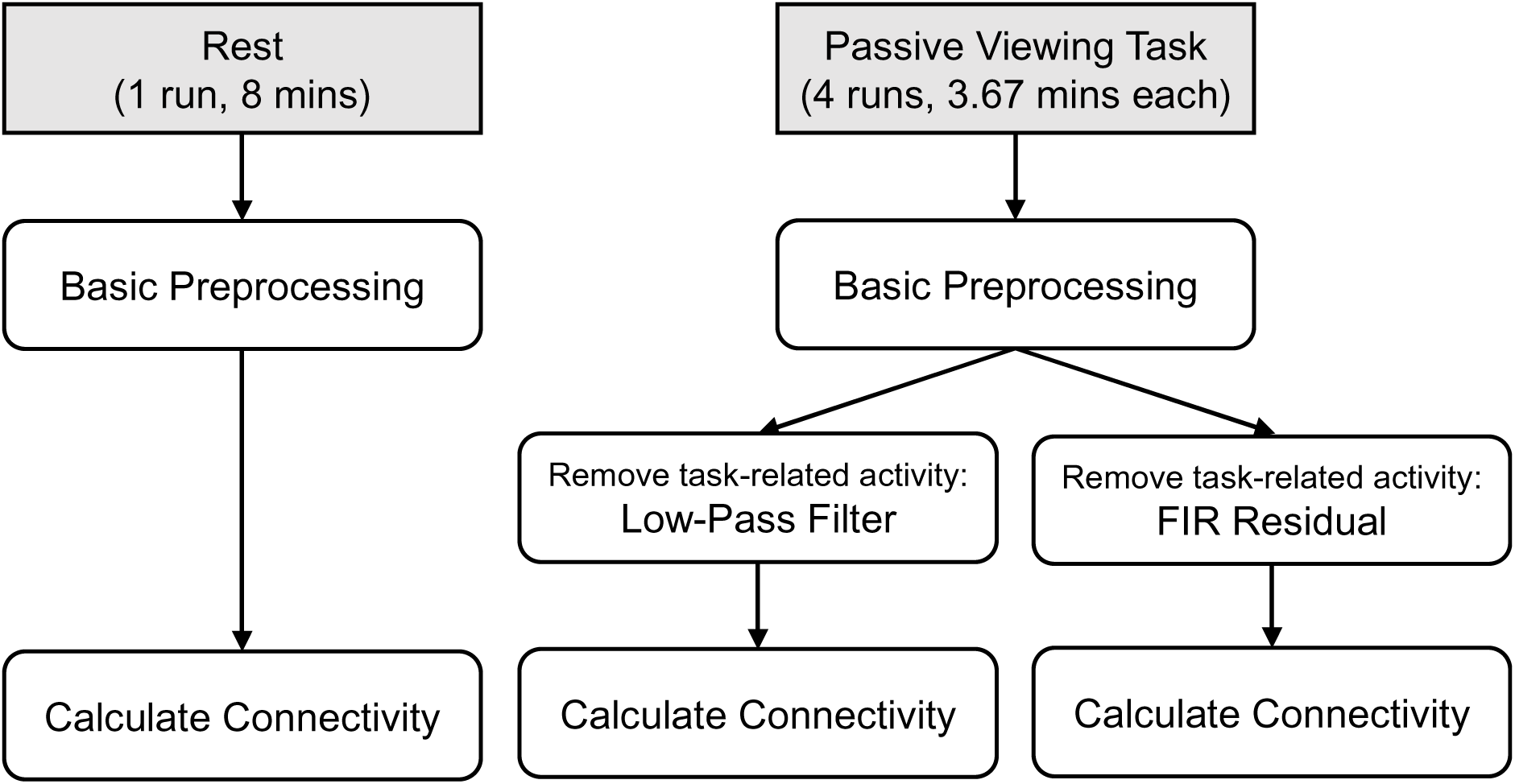
Overview of preprocessing and analysis pipeline. Resting-state and passive viewing data were subjected to the same preprocessing steps (see fMRI Preprocessing). ROI-to-ROI connectivity was measured from the preprocessed resting-state data, while data from the passive viewing task were subjected to an additional step of preprocessing. Task-related activity was removed from the passive viewing data using the low-pass filter (LPF) and FIR residuals (FIR) approach. Connectivity was calculated from the LPF data and from the FIR data, resulting in two sets of task-based connections that were then compared to the “gold standard” rest connectivity.

#### 2.2.4 fMRI Preprocessing

Dicom files were converted to nifti format using the “dcm2nii” function from MRIcron (https://www.nitrc.org/projects/mricron) and organized in BIDS format. fMRI preprocessing and data analysis were carried out using FEAT (fMRI Expert Analysis Tool), version 6.00, part of FSL (www.fmrib.ox.ac.uk/fsl) and custom scripts. All functional runs were brain extracted using BET and motion corrected within each run using McFlirt. The functional and MPRAGE anatomical scans of each subject were coregistered to their first functional volume by rigid/affine transformations using the Advanced Neuroimaging Tools (ANTs: http://stnava.github.io/ANTs/). Noise components from the co-registered functionals were identified and removed using Independent Component Analysis-based Automatic Removal of Motion Artifacts (ICA-AROMA) (Pruim et al., 2015; https://github.com/maartenmennes/ICA-AROMA). A high- pass temporal filter (100 s) was then applied to the denoised functionals. The resulting timeseries were then ready for the computation of connectivity measures (resting state fMRI) or for the removal of task-evoked activity (task-based fMRI).

#### 2.2.5 Removing Task-Evoked Activity from Task-Based fMRI

Functional timeseries from task-based fMRI underwent an additional processing step—removal of task-evoked activity—prior to calculating connectivity measures. Here, we utilized and compared two previously used methods proposed to remove task-evoked activity, low-pass filter (Frank, Bowman, et al., 2019; Frank, Preston, et al., 2019; Tambini et al., 2017) and FIR residuals (Al-Aidroos et al., 2012; Fair et al., 2007; Norman-Haignere et al., 2012). In the low-pass filtered method, we applied a low-pass filter with 16 s cutoff to remove frequencies in the BOLD signal that were at or higher than the 12 s task frequency. A conservative threshold of 16 s was used to ensure all task-related activity was removed. The filtered functional timeseries were then utilized for further connectivity analyses. We will refer to this low-pass filtered task-based dataset as “LPF”.

For the FIR residuals method, we ran a general linear model, modeling task-evoked BOLD signal using a finite impulse response (FIR) function rather than canonical hemodynamic response function to maximize model fits. The model included 6 FIR basis functions to estimate activity at each 2-s TR time point within the 12-s trial window (Glover, 1999; Kay et al., 2008). The model also included the nuisance regressors outlined by Power et al. (2012), including the 6 motion parameters, cerebrospinal fluid signal, white matter signal, whole brain signal and each of their derivatives. After regressing out the task-evoked activation, we obtained the residual timeseries (activation not accounted for by the task) to be utilized for further connectivity analyses. We will refer to this residual timeseries dataset as “FIR”. Thus, from the single original task-based fMRI dataset, we generated two datasets for measuring background connectivity, one using LPF and one using the FIR residual method to account for task-evoked activity.

#### 2.2.6 Measuring Functional Connectivity

As we were interested in the stability of overall connectivity patterns across task and rest, we focus on whole brain ROI-to-ROI connectivity. To assess the potential of each method to reproduce established functional brain networks, we utilized a brain atlas that contains information regarding network membership for each segmented brain region. We adopted the Schaefer et al. (2018) parcellation scheme, containing 100 parcels organized into 7 cortical networks (Yeo et al., 2011). This allowed us to compare 100x100 matrices of ROI-to-ROI connections (symmetrical along the diagonal) as well as summarize those connections at the network level.

The previous steps provided us with three timeseries datasets: resting-state fMRI (rest), low-pass filtered data from task-based fMRI (LPF), and FIR residuals from task-based fMRI (FIR). The same procedures for measuring connectivity were applied to each of the three datasets. Timeseries were first extracted from each of the 100 parcels. Volumes that exceeded FD > .5 mm or DVARS > .5% were “scrubbed” or removed from the timeseries (Power et al., 2012). Pairwise partial correlations were conducted between the scrubbed timeseries of the 100 parcels controlling for nuisance regressors, including the 6 motion parameters, cerebrospinal fluid signal, white matter signal, whole brain signal and each of their derivatives (Power et al., 2012). Note that although the nuisance regressors were already included in the FIR models used to obtain the residual timeseries, they were also partialled out when calculating connectivity to match the procedures with the rest and LPF datasets. The results did not change when using FIR residual background connectivity measured without the nuisance regressors.

Background connectivity estimates for the task-based (LPF and FIR residual) datasets were calculated separately within each run and then averaged across runs. Three individual runs of the passive viewing task were excluded for program errors (one participant), participant falling asleep (one participant), and non-compliance with study protocols (one participant). The connectivity values for these subjects thus reflects the average of three rather than four runs.

The above procedures generated three 100x100 ROI-to-ROI correlation matrices for each subject: a rest-based connectivity matrix (rest), a background connectivity matrix obtained after low-pass filtering (LPF), and a background connectivity matrix obtained from FIR residuals (FIR). These matrices were then utilized to compare rest and background connectivity in terms of (1) within-subject connectivity profiles, (2) potential to reproduce known network structures, and (3) individual differences in connectivity strength. Please note that the correlation coefficients denoting connectivity strength are reported raw in text and figures for intuitive reading but were always Fisher z transformed before being used for further statistical analyses, per standard recommendations (Dunn & Clark, 1969; Silver & Dunlap, 1987).

### 2.3. Analyses comparing rest and background connectivity

#### 2.3.1 Similarity of subject-specific connectivity profiles

It has been argued that an individual’s connectivity profile at rest contains idiosyncratic features that may serve as their “connectivity fingerprint” (Finn et al., 2015). To evaluate whether task-based functional connectivity can be similarly utilized, we asked how closely each individual subject’s background connectivity matrices (FIR, LPF) matched their rest-based connectivity matrix. To quantify the similarity for each subject, the upper triangles (not including the diagonal) of the subject-specific correlation matrices were vectorized and correlated between each method. As there were 100 ROIs in the atlas, there were 4,950 unique ROI-to-ROI correlation values for each participant and dataset (rest, LPF, FIR) that represented their subject-specific connectivity patterns. To evaluate how similar such subject-specific connectivity patterns are between task and rest, Spearman ranked correlations were conducted between rest connectivity patterns and patterns obtained from each of the background connectivity measures (rest x LPF and rest x FIR). The two background connectivity measures were also correlated with each other (LPF x FIR) to evaluate how similar the subject-specific estimates were between the two approaches to removing task-evoked signal when applied to the same original task-based dataset. To evaluate whether one task-based method (LPF or FIR) consistently produces patterns more similar to those obtained from rest, we compared the subject-specific rest x LPF similarity scores with rest x FIR similarity scores, using a paired-samples t-test. As noted above, the similarity scores (Spearman’s correlation coefficients) were Fisher z transformed prior to the statistical tests.

#### 2.3.2 Reproducing network structure of the brain

We next asked how well each of the background connectivity methods reproduce the 7 pre-defined resting-state cortical networks as implemented in the 7-network version of the Schaefer atlas (Schaefer et al., 2018; Yeo et al., 2011). For each participant, we calculated the average within-network and between-network connectivity produced by each method (rest, LPF, FIR). Within-network connectivity was calculated by averaging the estimates for all unique within-network ROI-to-ROI connections.

Between-network connectivity was calculated by averaging all unique between-network ROI-to-ROI connections. We then ran a 3 (method: rest, LPF, FIR) x 2 (connection type: within-network, between-network) repeated measures ANOVA to compare how defined functional networks were in each dataset. We expected within-network connectivity to be greater than between-network connectivity, consistent with the cortical network labels assigned to each parcel by Schaefer and colleagues (2018). Of main interest, however, was the interaction between method and the type of connection (within-network or between-network) to compare whether the network structure was more or less pronounced in any dataset.

#### 2.3.3 Stability of individual differences in connectivity

As the last analysis, we compared the stability of individual differences in connection strengths between rest and the background connectivity methods. In other words, we wondered whether individual differences (e.g., some subjects having a particularly strong connection between two regions) identified in rest also consistently appear in the background connectivity estimates. We addressed this question on the level of ROI-to-ROI connections and on the level of networks. For each ROI-to-ROI connection, we calculated the across-subject Spearman ranked correlations in the connectivity estimates between rest and LPF, rest and FIR, and LPF and FIR. For example, we took the ROI1-ROI2 connectivity estimates for all subjects from the rest data and correlated them with subjects’ ROI1-ROI2 connectivity estimates from the LPF data. To examine individual differences in large-scale networks, we additionally collapsed the 100x100 ROI-to-ROI connectivity estimates (excluding connections on the diagonal) from each subject to a 7x7 network-to-network matrix. For each unique network-to-network connection, we conducted across-subject Spearman ranked correlations in the connectivity estimates between rest and LPF, rest and FIR, and LPF and FIR. As always, the rank correlations were Fisher z transformed for analysis. In addition to assessing the overall similarity of rest and background connectivity-based estimates of individual differences, we also tested whether individual differences from rest connectivity are more consistent with individual differences in the LPF-based or FIR-based connectivity measures. A paired-samples t-test was conducted to compare the rest x LPF correlations to the rest x FIR correlations across all connections.

## 3. Results

### 3.1 Similarity of subject-specific connectivity profiles across task and rest

We first tested how similar subject-specific patterns of ROI-to-ROI connectivity were between rest and each of the background connectivity methods. For each participant, we correlated their ROI-to-ROI connectivity pattern from rest with the background connectivity measures from the LPF dataset (rest x LPF) and from the FIR dataset (rest x FIR). The connectivity matrices, averaged across participants, are presented in **Figure 2**. The subject-specific background connectivity patterns estimated from the task-based fMRI were moderately to highly correlated with the same participant’s connectivity patterns estimated from rest (Median rest x LPF pattern similarity rho = 0.63; Median rest x FIR pattern similarity rho = 0.74). Rest connectivity patterns were more closely matched by background connectivity patterns derived using the FIR method (*Mean rho(z)* = 0.93, *SD* = 0.19) than LPF method (*Mean rho(z)* = 0.75, *SD* = 0.18; *t*(55) = -18.95, *p* < 0.001, *ηg* = 0.87; **Figure 3A**). Nevertheless, patterns of background connectivity from LPF and FIR residuals data were highly correlated with each other (Median rho = 0.85) and thus both methods likely produce very similar results in practice.

**Figure 2.**
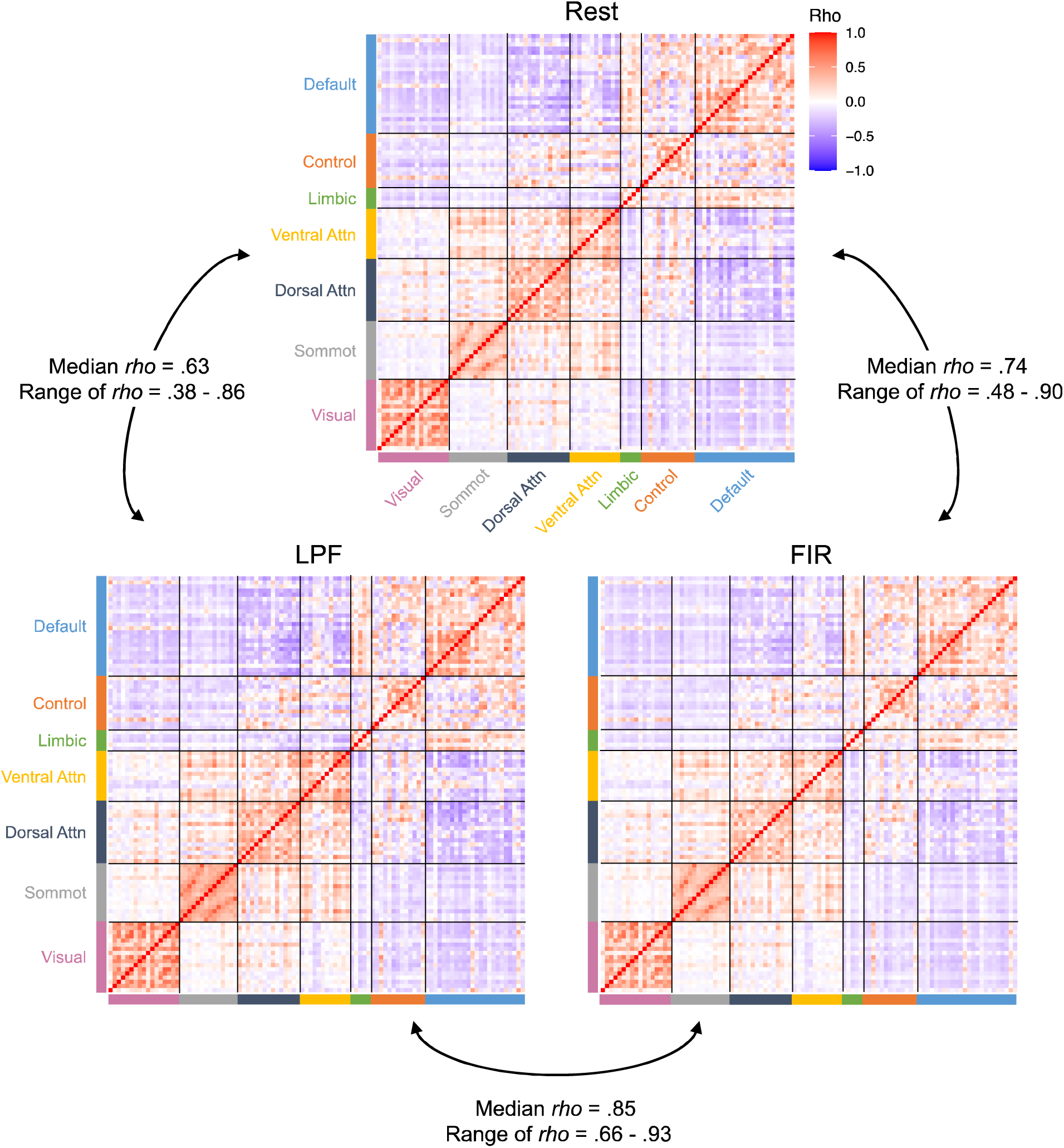
Group averaged ROI-to-ROI correlation matrices for each method. The ROI-to-ROI correlation matrices are shown for the resting-state data, task-based LPF data, and the task-based FIR data. The selected ROIs are from the Schaefer et al. (2018) parcellation scheme and are organized into 7 cortical networks (Default, Frontoparietal Control, Limbic, Ventral Attention, Dorsal Attention, Somatomotor, Visual; Yeo et al., 2011). The background connectivity matrices displayed are averaged across the four runs of passive viewing. For ease of interpretation, the matrices display the raw correlations prior to Fisher z transformation. For each subject, pairwise correlations were conducted between the three matrices to determine how well each background connectivity method reproduces the given individual’s pattern of connections found during rest. The median and range of these within-subject correlations are displayed.

**Figure 3.**
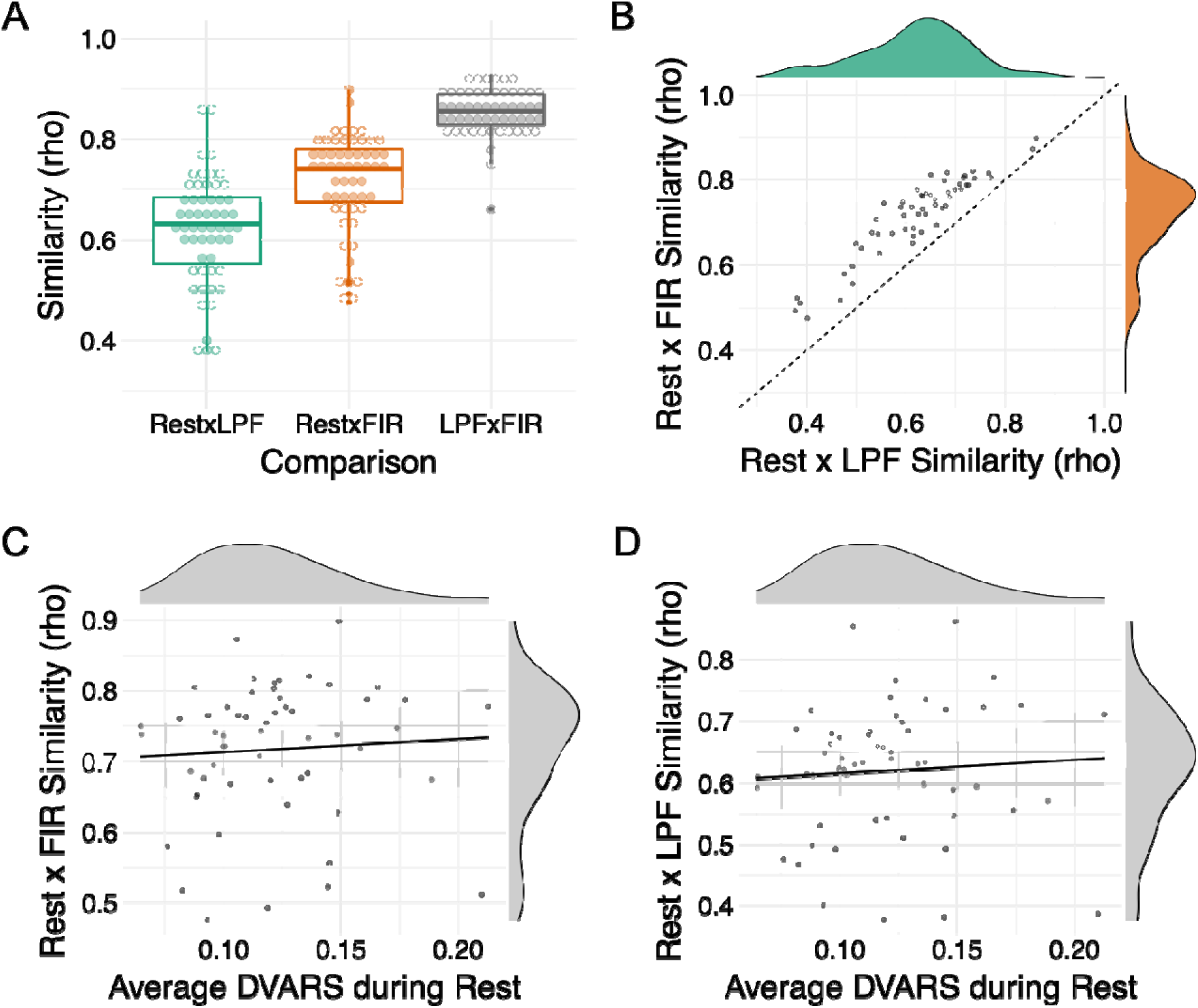
Similarity in individual patterns of connectivity. For each subject, pairwise correlations were conducted to assess the similarity between the individual patterns of connections produced by each method. For ease of interpretation, raw correlations prior to Fisher z transformation are displayed. **A.** The distribution of within-subject similarity scores for each comparison. Each dot represents a participant. The box represents the interquartile range (Q1-Q3), the middle bar represents median (Q2), and the whiskers represent the minimum and maximum values. Points outside of the whisker (> 1.5 * interquartile range) are defined as outliers. All pairwise comparisons are statistically significant at *p* < 0.001 (rest-LPF < rest-FIR < LPF-FIR). **B.** A scatter plot showing a strong correlation between rest-LPF similarity scores and rest-FIR similarity scores (*r* = 0.93, *p* < 0.001). Those who showed low similarity between rest and task connectivity patterns did so irrespective of the background connectivity method (LPF vs. FIR residuals). Note that all subjects showed higher rest-FIR similarity than rest-LPF similarity (shown by all dots above the line x=y). **C, D.** Scatter plots showing the correlation between average DVARS during rest and subject-specific similarity scores between rest and task (C: rest x FIR similarity, D: rest x LPF similarity). The correlations were numerically positive but not statistically significant. Thus, low similarity in connectivity estimates between rest and task in some subjects was not clearly attributable to motion.

Although the subject-specific patterns of connectivity were typically consistent between task and rest, some subjects exhibited consistency as low as rho near 0.4 (see ranges presented in **Figure 2** and the distributions of similarity scores in **Figure 3A**).

We were interested if those who demonstrate low rest-LPF similarity also show low rest- FIR similarity. Indeed, the pattern similarity scores were highly correlated (*r* = 0.93, *p* < 0.001; **Figure 3B**). This indicates that low consistency scores were not driven by a specific method of calculating background connectivity but instead were found for a given individual with either method.

We reasoned that perhaps lower similarity may be seen for participants who moved more during scanning and whose connectivity estimates may thus be less reliable. To test this idea, we correlated the rest x task pattern similarity scores with four indices of individual differences in motion: average DVARS, average FD, max DVARS, and max FD. The results did *not* show a clear relationship to motion. For rest x FIR similarity scores, the only numerical relationship we found was with the average DVARS during the rest scan, such that subjects who had higher rest x task similarity tended to have higher average DVARS (*rho*(54) = 0.19, *p* = 0.166) (**Figure 3C**). The rest x LPF similarity scores showed a similar relationship (*rho*(54) = 0.18, *p* = 0.182). However, the relationships were relatively weak and no correlations with motion reached significance (all uncorrected *p* > .05). Thus, while motion may play a role in the consistency of connectivity patterns between task and rest, our data do not allow us to clearly attribute lower task-rest consistency in some subjects to motion.

### 3.2 Reproducing network structure of the brain

Next, we asked how well each of the background connectivity methods reproduces large-scale brain networks (Schaefer et al., 2018; Yeo et al., 2011), using the network definitions from the Schaefer et al. (2018) parcellation atlas. A visual inspection of Figure 2 indicates that network structure was clearly observable in all three datasets (rest, LPF, FIR). To formally quantify the network organization, all the unique connections from the 100 x 100 ROI-to-ROI connection matrices (excluding the diagonal) were divided into within-network connections and between-network connections, separately for each subject. The average within-network and average between-network connection values from each subject and dataset were then submitted to a 3 (rest, LPF, FIR) x 2 (within-network, between-network) repeated measures ANOVA to test whether the strength of network organization varies among datasets (**Figure 4**).

**Figure 4.**
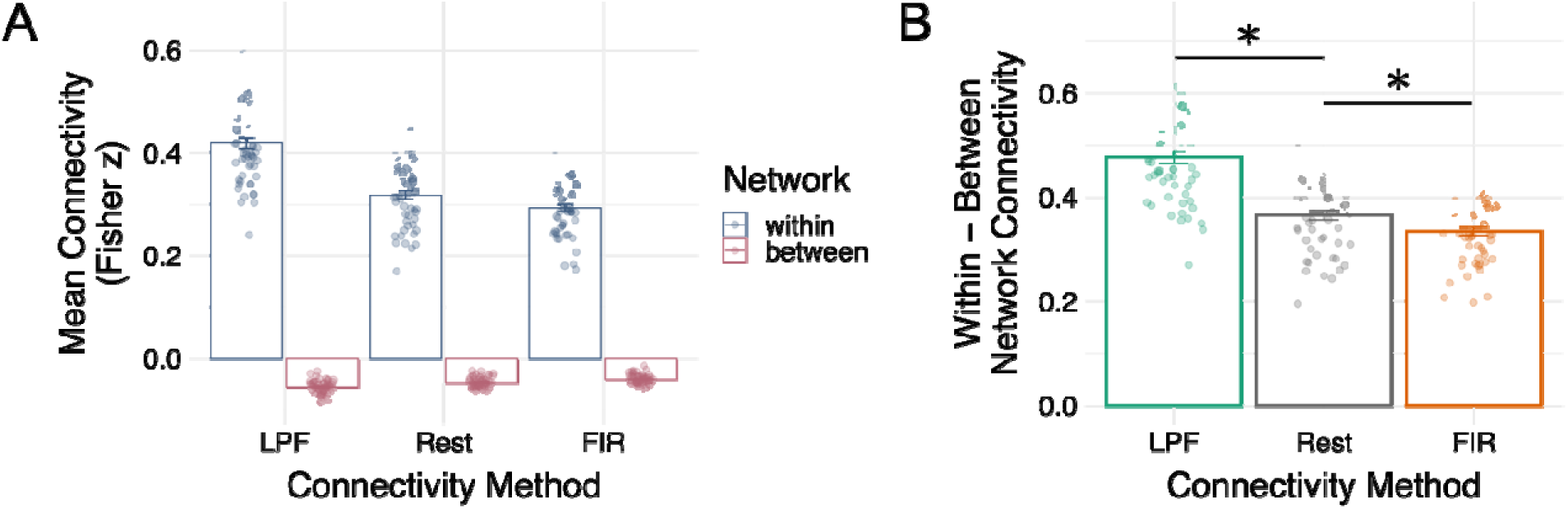
Reproducing resting-state network structure. **A.** Mean within-network and between-network connectivity estimates for each method are plotted with data points for individual subjects. Error bars denote +/- 1 SE. **B.** The mean within-network minus between-network connectivity differences, plotted for each method. Data points for individual subjects are included. Error bars denote +/- 1 SE.

As expected, average connectivity within networks (*Mean rho(z)* = 0.34, *SD* = 0.09) was significantly higher than between networks (*Mean rho(z)* = -0.05, *SD* = 0.01; *F*(1,55) = 2125.79, *p* < 0.001, *η^2^g* = 0.95), confirming that the network organization of the Schaefer et al. (2018) parcellation was reflected in our data. We also found a main effect of connectivity method (*F*(1.41,77.6) = 135.13, *p* < 0.001, *η^2^g* = 0.21, Greenhouse Geiser corrected). Follow-up comparisons revealed the LPF method (*Mean rho(z)* = 0.18, *SD* = 0.25) produced significantly higher connections compared to rest (*Mean rho(z)* = 0.135, *SD* = 0.19; *t*(55) = 9.97, *p* < 0.001, *η^2^g* = 0.36) and FIR methods (*Mean rho(z)* = 0.125, *SD* = 0.17; *t*(55) = 19.24, *p* < 0.001, *η^2^g* = 0.48), while the FIR method produced the smallest average connectivity values (comparison to rest: *t*(55) = 3.02, *p* = 0.004, *η^2^g* = 0.04).

Of main interest, however, was the interaction between the connectivity method (rest, LPF, FIR) and the type of connection (within-network, between-network). We found a significant interaction indicating that the network structure (the difference between within-network and between-network connectivity) varied among datasets (*F*(1.41,77.6) = 123.29, *p* < 0.001, *ηg2* = 0.30, Greenhouse Geiser corrected; **Figure 4**). Interestingly, network structure was most pronounced in the LPF-derived background connectivity estimates, followed by rest, followed by FIR residual-derived background connectivity estimates (**Figure 4B**). Follow-up pairwise t-tests were all statistically significant: LPF compared to rest (*t*(55) = 9.01, *p* < 0.001, *η^2^g* = 0.32), rest vs. FIR (*t*(55) = 3.86, *p* < 0.001, *η^2^g* = 0.05), and LPF vs. FIR (*t*(55) = 18.65, *p* < 0.001, *η^2^g* = .46). The smallest difference was between rest and the FIR background connectivity method rather than between the two background connectivity methods, with FIR method producing somewhat less pronounced network structure (see also **Figure 2** for the full ROI-to-ROI matrices further demonstrating this effect). Nevertheless, network structure was clearly reflected in all three datasets, with significantly higher within-network than between network connectivity in every dataset and every individual subject (**Figure 4**).

### 3.3 Stability of individual differences in connectivity across task and rest

Prior work suggested that individual differences in resting-state connectivity are meaningful, related to various individual characteristics, such as personality, cognition, or clinical symptoms (Finn et al., 2015; Liu et al., 2019; Reinen et al., 2018; Rosenberg et al., 2015; Takamura & Hanakawa, 2017; Toschi et al., 2018). Here, we asked whether individual differences in background connectivity are similar to those observed at rest. First, we examined the stability of individual differences on the level of ROI-to-ROI connections. For each ROI-to-ROI connection, across-subject correlations were conducted between rest and LPF, rest and FIR, and LPF and FIR. For example, subject estimates in ROI1-ROI2 connectivity during rest were correlated with subject estimates in ROI1-ROI2 connectivity derived from the LPF method. This resulted in an estimate of stability for each of the 4,950 ROI-to-ROI connections for each of the three comparisons (rest x LPF, rest x FIR, and LPF x FIR). The between-datasets similarity of individual differences estimates for each connection are visualized in **Figure 5** and summarized in **Table 1** and **Figure 6**.

**Figure 5.**
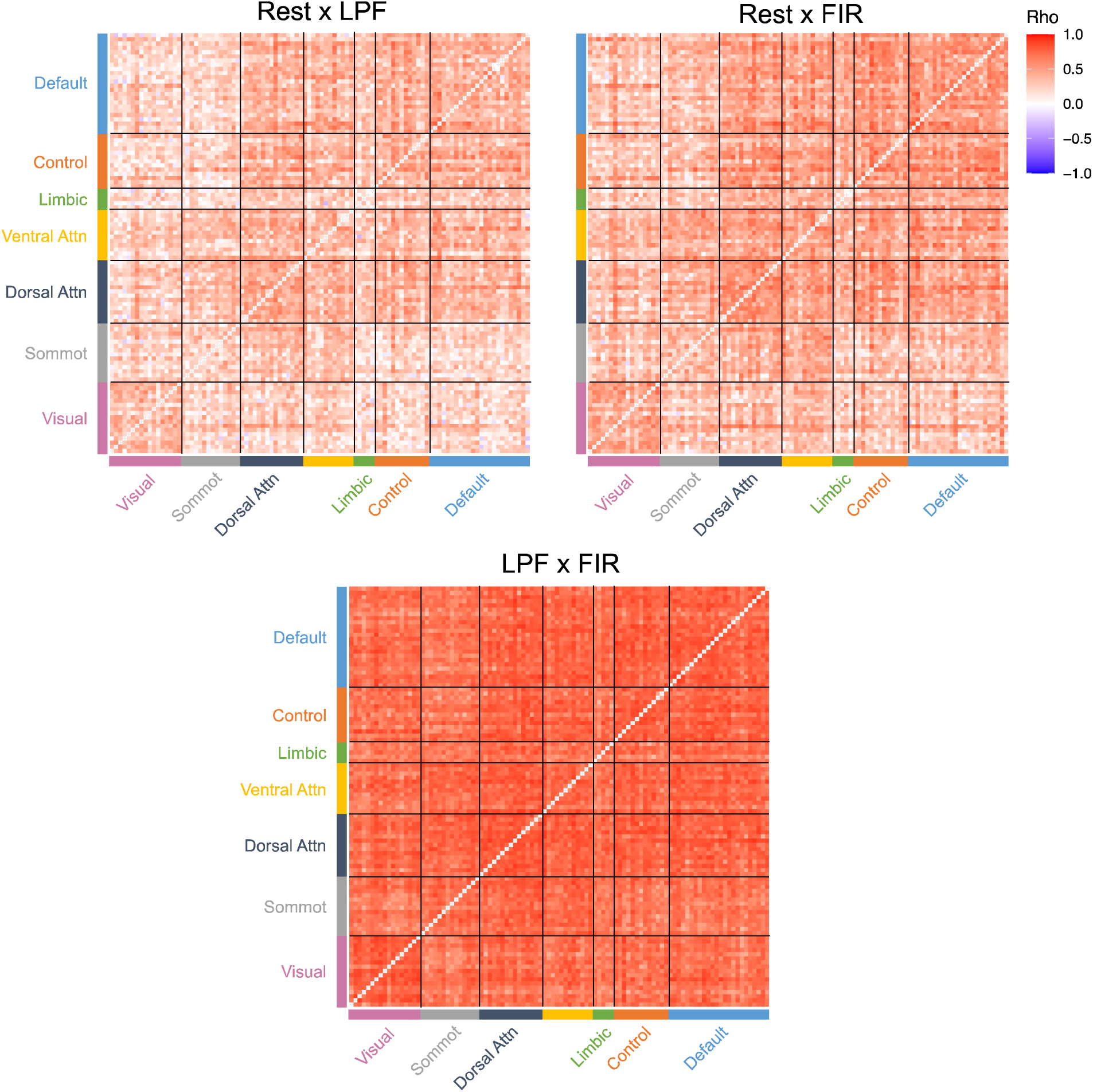
Stability of individual differences for each ROI-to-ROI connection. For each unique ROI-to-ROI connection, individual differences in connectivity estimates were correlated between rest and LPF, rest and FIR, and LPF and FIR datasets. For ease of interpretation, the raw correlations prior to Fisher z transformation are shown.

**Figure 6.**
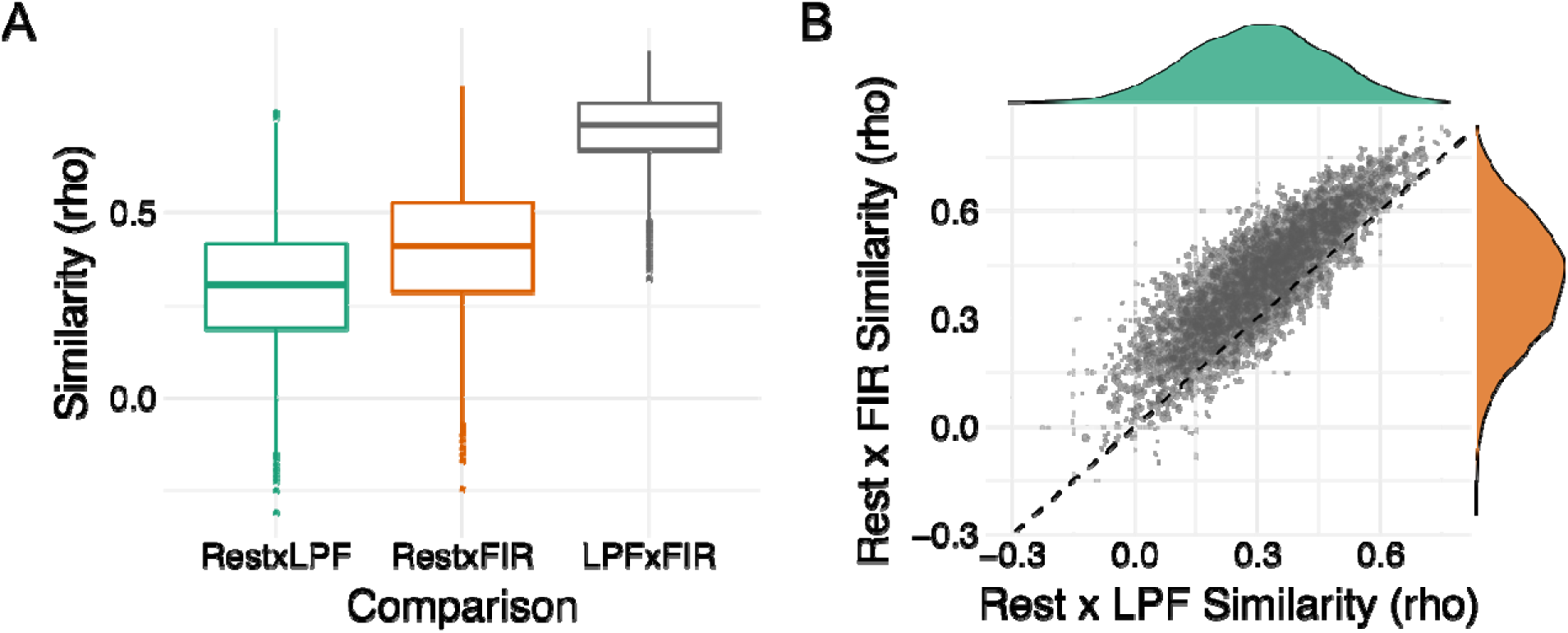
Stability of individual differences between the methods. For each ROI-to-ROI connection, individual differences in connectivity were calculated between two methods (rest x LPF, rest x FIR, and LPF x FIR). For ease of interpretation, raw correlations prior to Fisher z transformation are displayed. **A.** The boxplot of similarity scores (Spearman rho) for all unique ROI-to-ROI connections. The box represents the interquartile range (Q1-Q3), the middle bar represents median (Q2), and the whiskers represent the minimum and maximum values. All pairwise comparisons are statistically significant at *p* < 0.001 (rest-LPF < rest-FIR < LPF-FIR). **B.** Scatterplot showing a strong correlation between rest-LPF similarity and rest-FIR similarity (*r* = 0.85, *p* < .001). The connections that showed low similarity between rest and FIR also showed low similarity between rest and LPF, suggesting that some connections showed relatively low stability of individual differences between rest and task that was not driven by a particular method for removing task-related activity. Dashed line indicates where x = y.

**Table 1.**
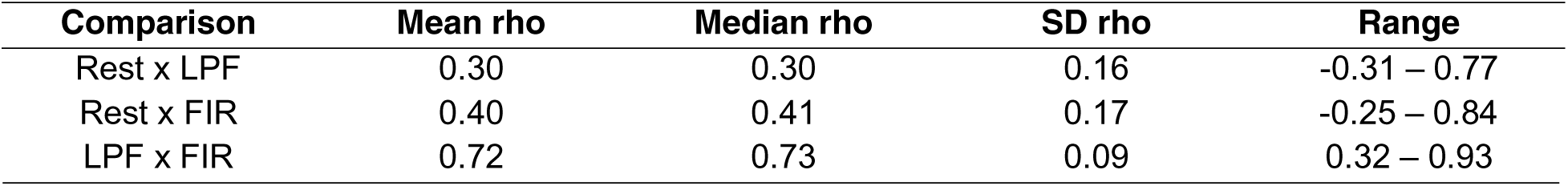
Summary of correlations in individual differences for ROI-to-ROI connections Comparison Mean rho Median rho SD rho Range

On average, individual differences in background connectivity were weakly to moderately correlated with individual differences in resting-state connectivity, suggesting that individual differences in specific ROI-to-ROI connections are not always stable from rest to task (**Table 1, Figure 6A**). Individual differences in background connectivity obtained from the FIR method were more correlated with rest than were individual differences obtained from the LPF method (*t*(4949) = 78.22, *p* < 0.001, *η^2^g* = 0.09).

Individual differences for some connections showed especially low similarity between rest and task (lowest *rho* = -0.31). To determine whether rest-task differences were common for certain connections or whether they were driven by a particular background connectivity method, we correlated rest x LPF similarity scores with rest x FIR similarity scores. We found that rest-FIR similarity was strongly and positively correlated with rest-LPF similarity (*rho*(4948) = 0.85, *p* < 0.001), with connections showing low similarity in individual differences between rest and LPF also showing low similarity between rest and FIR (**Figure 6B**). This finding suggests that for some regions, individual differences in connection strength can vary considerably between rest and task, irrespective of the method used for removing task-evoked activity (LPF, FIR).

Notably, given research on task-driven changes in connectivity (Cole et al., 2021; Elton & Gao, 2015; Rissman et al., 2004), these differences between task and rest may be meaningful, reflecting differential modulation of regional or network connections in different subjects. We thus wanted to explore the degree of stability across task and rest for different connections more qualitatively. First, we collapsed across the 100x100 ROI-to-ROI connections to 7x7 network-to-network connections, reducing the number of connections to consider while respecting known functional organization of the brain.

Individual differences in connectivity for each of the unique network-to-network connections were then correlated between rest and LPF, rest and FIR, and LPF and FIR. The correlations, visualized in **Figure 7** and summarized in **Table 2**, are of similar average strength as those found on the level of ROI-to-ROI connections. Similar to the ROI-to-ROI findings, individual differences in network-to-network connections demonstrated greater rest-FIR similarity than rest-LPF similarity (*t*(27) = 7.93, *p* < 0.001, *ηg2* = 0.21).

**Figure 7.**
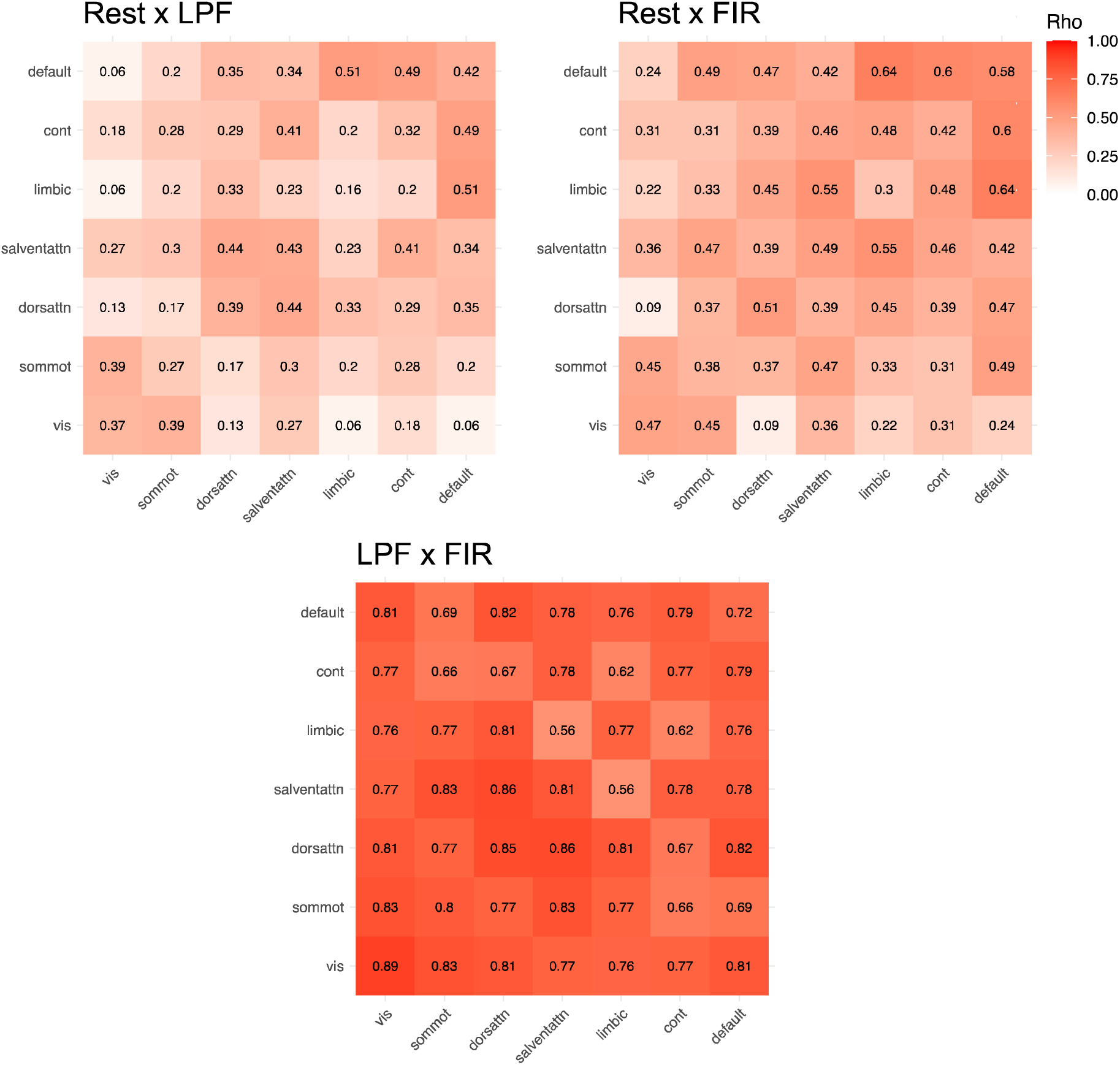
Stability of individual differences in network-to-network connections. The 100x100 ROI-to-ROI connections were collapsed into 7x7 network-to-network connections. For each network-to-network connection, individual differences in connectivity were correlated between two methods (rest x LPF, rest x FIR, LPF x FIR). For ease of interpretation, the correlations (rho) prior to Fisher z transformation are reported.

**Table 2.**
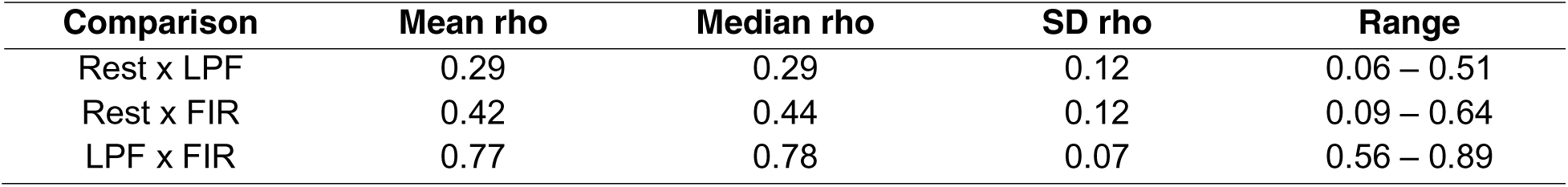
Summary of correlations in individual differences for network-to-network connections

| Comparison | Mean rho | Median rho | SD rho | Range |
| --- | --- | --- | --- | --- |
| Rest x LPF | 0.29 | 0.29 | 0.12 | 0.06 – 0.51 |
| Rest x FIR | 0.42 | 0.44 | 0.12 | 0.09 – 0.64 |
| LPF x FIR | 0.77 | 0.78 | 0.07 | 0.56 – 0.89 |

As can be observed from Figure 7, individual differences in the default mode network connections appeared the most similar between rest and task, both within the default network and between the default network and other networks. Individual differences of visual network connections, especially with the dorsal attention network and perhaps limbic and default networks (for LPF), appeared the least similar. Thus, individual differences in connectivity were not always stable from rest to task, driven by some connections more than others. In contrast, individual differences estimates of background connectivity were fairly consistent across the two analytical methods (LPF, FIR), both at the level of individual connections (**Table 1, Figure 5, 6**) and at the level of network connections (**Table 2, Figure 7**).

## 4. Discussion

Resting-state functionality connectivity is widely used for measuring functional interactions across the brain, substantially contributing to our understanding of the brain organization common across individuals. Furthermore, connectivity studies have also revealed the existence of individual differences in connectivity strengths and patterns, which have been linked to behavioral, clinical, cognitive, age, and other individual factors (Ferreira & Busatto, 2013; Fong et al., 2019; Geerligs et al., 2015; Liu et al., 2019; Poole et al., 2016; Reinen et al., 2018; Tracy & Doucet, 2015). Previous research suggests that brain activity occurring in the background of a task can be used to reproduce resting-state-like connectivity profiles (Cole et al., 2014; Gratton et al., 2018; Kraus et al., 2021), allowing researchers to get further use out of existing task-based fMRI datasets. Here, we tested this idea along with two methods for removing task-related activity in a slow event-related fMRI task design: low-pass filter and general linear model. Our results demonstrated that background connectivity derived from task-based fMRI successfully reproduced large-scale cortical networks found at rest and largely maintained within-subject patterns of ROI-to-ROI connectivity. In contrast, individual differences in connectivity strength were less stable across task and rest for many connections, potentially reflecting task-driven modulations of connectivity strength that differ across individuals. Across all analyses, rest-based connectivity measures were more closely approximated by background connectivity derived after general linear modeling of task-evoked activation than low-pass filtering of task-evoked activation.

Nevertheless, both methods for removing task-evoked activity generated connectivity patterns similar to rest and to one another, suggesting both are viable approaches for measuring background connectivity.

A significant contribution of resting-state connectivity has been the identification of large-scale networks, which can be leveraged for clinical research (Reinen et al., 2018; Takamura & Hanakawa, 2017; Tracy & Doucet, 2015) and understanding human behavior and cognition (Cohen & D’Esposito, 2016; Shine et al., 2016). Here, we reproduced large scale cortical networks during rest (Schaefer et al., 2018; Yeo et al., 2011) and demonstrated that functional networks are also maintained in background connectivity derived from task-based fMRI. Previous studies have shown similar organization of functional networks between rest and task (Bzdok et al., 2016; Smith et al., 2009), with tasks recruiting the same intrinsic networks derived from rest (Gess et al., 2014; Shah et al., 2016). Together, these findings suggest that large scale cortical networks initially identified from resting-state fMRI may reflect an inherent and stable organization of the brain.

One inspiration for our study was prior work suggesting that individuals display unique functional connectivity profiles that can be treated as a type of “connectivity fingerprint” (Finn et al., 2015). Moreover, the relative strength of individual connectivity has been shown to contain information about factors, such as mental health conditions, disease, cognitive abilities, and personality traits (Fong et al., 2019; Liu et al., 2019; Rosenberg et al., 2015; Takamura & Hanakawa, 2017; Toschi et al., 2018; Tracy & Doucet, 2015). Thus, we conducted two analyses focusing on individuals: idiosyncratic (within-subject) connectivity patterns and individual differences in connectivity strength. Aligned with the fingerprint hypothesis, we found that idiosyncratic patterns of ROI-to-ROI connectivity during task-based fMRI were highly similar to those found at rest for most participants. These results contribute to a growing literature that shows how connectivity profiles during a task are largely made up of idiosyncratic features (Cole et al., 2014; Gratton et al., 2018; Kraus et al., 2021).

As individual behaviors or traits can be linked to specific connections or networks, rather than a whole-brain profile (Gerraty et al., 2014; Klumpp et al., 2014; Poole et al., 2016; Qin et al., 2014; Toschi et al., 2018), we then asked whether individual differences for specific connections are maintained between rest and task-based fMRI. Unexpectedly, individual differences in ROI-to-ROI and network-to-network connections were only weakly to moderately correlated between rest and task. These differences are unlikely to reflect just noise, given that they were observed even when averaging across many connections for network-to-network analyses. Instead, the differences between rest and background connectivity may be driven by engagement of specific connections during the task. Indeed, Fair and colleagues (2007) showed that connections that were unique to the task-based data overlapped with regions that were also activated during the task. Thus, it is important to keep in mind that while background connectivity may contain information about resting-state-like connectivity profiles, they may also contain information regarding task-related behavior.

Our finding that individual differences in connectivity strength observed during rest are not always consistent with those observed during task should not be taken as an argument against using background connectivity measures in studies of individual differences. Prior studies have demonstrated the importance of task-modulated functional connectivity in predicting behavior (Elton & Gao, 2015; Greene et al., 2018; Jiang et al., 2020), with some suggesting it may even serve as a better predictor compared to rest (Greene et al., 2018; Jiang et al., 2020). Indeed, background connectivity has been shown to track different brain states, such as attentional states (Al-Aidroos et al., 2012), emotional states (Tambini et al., 2017), and encoding/retrieval states (Cooper & Ritchey, 2019). Moreover, Gratton and colleagues (2018) found that task modulations of functional networks were largely due to individual variations in connectivity changes between task and rest, which could reflect individual differences in task engagement. Here, we did not have a suitable behavioral measure of individual differences and our sample size would not provide sufficient power to compare brain-behavior relationship between task and rest. However, it would be informative to compare the functional significance of rest-derived and task-derived individual differences measures in future studies.

In addition to testing how rest-derived metrics compared to background connectivity metrics, we were interested in comparing two methods of removing task-related activation from slow event-related task-based designs: low-pass filtering and general linear model. When examining individuals, background connectivity derived from the FIR residual dataset demonstrated greater similarity with resting-state connectivity compared to the LPF dataset. This was true for both idiosyncratic patterns of connectivity and individual differences in the strength of connections. Interestingly, though there were differences in how the two methods compared to rest, our findings also suggested that background connectivity measures are highly similar between the two methods. Notably, the measures derived from the two methods of removing task-evoked activity were highly correlated with each other and were similarly related to rest-based measures. For example, the degree to which an individual’s connectivity profile matched between task and rest was similar for low-pass filtered data and FIR residuals. The degree to which individual differences were stable or changed across task and rest for a particular connection was also relatively consistent across the two background connectivity methods. Thus, any differences between task and rest did not seem driven by a particular method. While the FIR residuals method generates measures that more closely match rest-based measures, the LPF method provides nearly identical results while being computationally simpler and faster. Thus, researchers can choose either method of removing task-related activity based on their needs, with existing and future studies using one or the other being directly comparable. The only notable difference between the two background connectivity methods were found in the reproduction of the large-scale cortical networks, where the LPF method produced networks more pronounced than those found during rest while FIR method produced networks less pronounced than rest. Because the underlying network modularity must be the same between LPF and FIR dataset, as both are coming from the same raw data, these results indicate that the absolute value of connection strength is differentially affected by various analysis steps (such as signal smoothing) and may not be directly comparable unless identical analysis steps are used.

Here, we contribute new evidence that background connectivity can capture the stable and intrinsic structure of functional connectivity as found during rest. Specifically, we show how removing task-related activity can preserve the organization of large-scale functional networks, as well as idiosyncratic connectivity patterns within subjects. While idiosyncratic patterns are maintained, individual differences can vary between rest and task. How changes in individual differences from rest to task relates to behavior is still unknown and should be explored in future studies. Two previously utilized methods for removing task-related activity in a slow event-related fMRI design, applying a low-pass filter and extracting residuals from an FIR model, both appear to be viable approaches with different advantages. To our knowledge, this is the first study comparing the two methods for removing task-related activity (FIR residual and low-pass filtering), particularly in a slow-event related design. How these methods compare in blocked task-based fMRI designs is an open area for exploration in future studies. Together, our findings highlight the relevance of background connectivity in the “connectivity fingerprint” hypothesis and open the door for the further utilization of task-based fMRI datasets.

## Notes

This work was supported by the National Institute of Neurological Disorders and Stroke Grant R01-NS-112366 (to D.Z.).

The authors have no known conflicts of interest to disclose. The data for this study are available through OpenNeuro at https://openneuro.org/datasets/ds004349.

### Competing Interest Statement

The authors have declared no competing interest.

